# TipQAD: an automated tool for quantifying apical fluorescence dynamics in tip-growing cells

**DOI:** 10.1101/2020.03.01.960591

**Authors:** Asongu L. Tambo, Bir Bhanu, Nan Luo, Duoyan Rong, Fei Wang, Christian Craddock, Irene Lavagi, Zhenbiao Yang

## Abstract

Cell polarity is a fundamental property essential for the function and development of all cellular organisms. Tip growth is an extreme form of polar growth requiring spatiotemporally dynamic but highly coordinated cellular activities. Quantification of these dynamic activities is important for the systematic study of the mechanisms controlling cell polarity, but an automated and unbiased method for analyzing and quantification of tip-growing cells has been missing. In this paper, we developed a computational model and an associated tool called TipQAD for quantifying the spatiotemporal dynamics of fluorescence protein (FP) labeled molecules or structures in tip-growing cells. This tool robustly and accurately measured the spatial distribution and temporal dynamics of FP-labeled proteins in the cytoplasm or the plasma membrane at the tip region of several types of tip-growing cells. We also demonstrated its ability to analyze data from FRAP experiments. In conclusion, this tool showed great potential for automated measurements of the spatiotemporal dynamics of fluorescent-labeled cellular molecules and structures.

## Introduction

Cell polarity describes the asymmetric property of a cell in all cellular organisms and is critical for cell function and development. It is required for cell movement, morphogenesis, differentiation, polar and directional growth, and so on. All forms of polarity are achieved through the spatiotemporal dynamic networks of regulatory molecules and cellular structures. As such, there is a need for automated tools that can quickly and accurately quantify the dynamics of these cellular molecules and structures in time and space. Tip growth, also known as apical cell growth, is an extreme form of polar growth characterized by a unidirectional expansion from a particular polarized region of the cell and is frequently observed in many cell types such as pollen tubes, root hairs^1^, fungal hyphae, and animal neurons. The networks of signal or structural molecules for regulating the polarity and building the cell are highly dynamic and coordinated during the polar growth, as the nascent cell membrane and cell wall materials are inserted into the same region where the networks are localized. Hence tip-growing cells provide an excellent system for the study of spatiotemporally dynamic cellular activities.

Among model systems for studying the molecular/cellular network underlying tip growth, pollen tube is one of the most studied because it can be conveniently manipulated and imaged *in vitro* and because of various genetic and molecular tools available for this system. As in other walled tip-growing cell types, the growth of pollen tubes at the apex requires the local expansion of the cell wall driven by turgor pressure, while the mechanical properties of the cell wall determine the morphology of the cell^2–5^. The precise control of the deposition of cell surface materials is regulated by a highly coordinated signaling network, involving vesicular trafficking, cytoskeletal dynamic/re-organization, ROP (Rho-like GTPase from plants) signaling and fluxes of ions such as calcium^3,6,7^. This network is characterized by its remarkable spatiotemporal dynamics. Spatially, it is highly polarized: dense populations of exocytic vesicles, supported by longitudinal actin cables and an apical actin network, move towards the extreme apex of pollen tubes in a “reverse fountain” pattern, delivering cell membrane and cell wall components to the tip^7,8^, while endocytosis retrieves excessive materials from the subapical or apical regions^9^. Active ROP1 and its signaling network, localized to the cell tip, has been shown to play an essential role in maintaining the cell polarity by regulating actin filaments and exocytosis^10–12^. Calcium ions also show a tip-high gradient and are crucial for pollen tube growth by participating in multiple pathways including the ROP signaling^13^. Temporally, this system is oscillatory and extremely dynamic: ROP1 activity, calcium concentration, and accumulation of vesicles at the cell tip exhibit rapid and correlated oscillations resulting in oscillating growth^14–16^. The mechanisms underlying the polar growth of other tip-growing cells, such as root hair, fungal hyphae, and neurons, are evolutionarily conserved. Therefore, the study of polar growth in these cell models relies heavily upon the analysis of the distribution and dynamics of various molecules and structures, yielding large amounts of video image data that require quantification and interpretation, which is extremely time-consuming by manual measurements.

In the current study, we develop TipQAD (Tip Quantification of Apical Dynamics), which is an automated tool for analyzing the spatial distribution and temporal dynamics of fluorescently labeled molecules/structures on the plasma membrane (PM) and in the cytoplasm at the apex of tip-growing cells. The proposed method is novel for the following reasons: (1) tip detection is done by optimizing intuitive functions based on the expectations of the geometry of tip-growing cells^4,6,17^; (2) a fast, fully automated measurement method is developed to replace the manual measurements that are currently made in the study of tip growth. Given a video frame, the method performs image segmentation (using user feedback on the first image) to obtain the shape of the cell. Then tip detection is used to determine the location and vertex of the tip region. Based on user-defined distances, measurements are made along the cell membrane and in the cytoplasm if required. Here we used this tool to analyze the spatiotemporal dynamics of fluorescence protein (FP)-tagged proteins localized in the cytoplasm in *Arabidopsis* pollen tubes, Arabidopsis root hairs. The accuracy of our automated program was confirmed by comparing the results with manual measurements.

## Results

### Computer vision techniques mediated recognition of tip-growing cells

Our computational model assumes that a tip-growing cell is made up of a cylindrical base and a hemispherical tip^4,6,17^ (Figure 1). After segregating the tip-growing cell, our model focuses on determining the best location for the apex (vertex) of the tip-growing cell in a given image(s). To ensure continuity in detecting the cell apical region, we assume that the apical region of the cell in the current frame is close to that of the previous frame. As such, we can limit the search space for the apical region in the current frame to the contour locations that are next to the apical region in the previous frame. Finally, our method assumes that the frames to be analyzed contain a single pollen tube.

**Figure 1:**
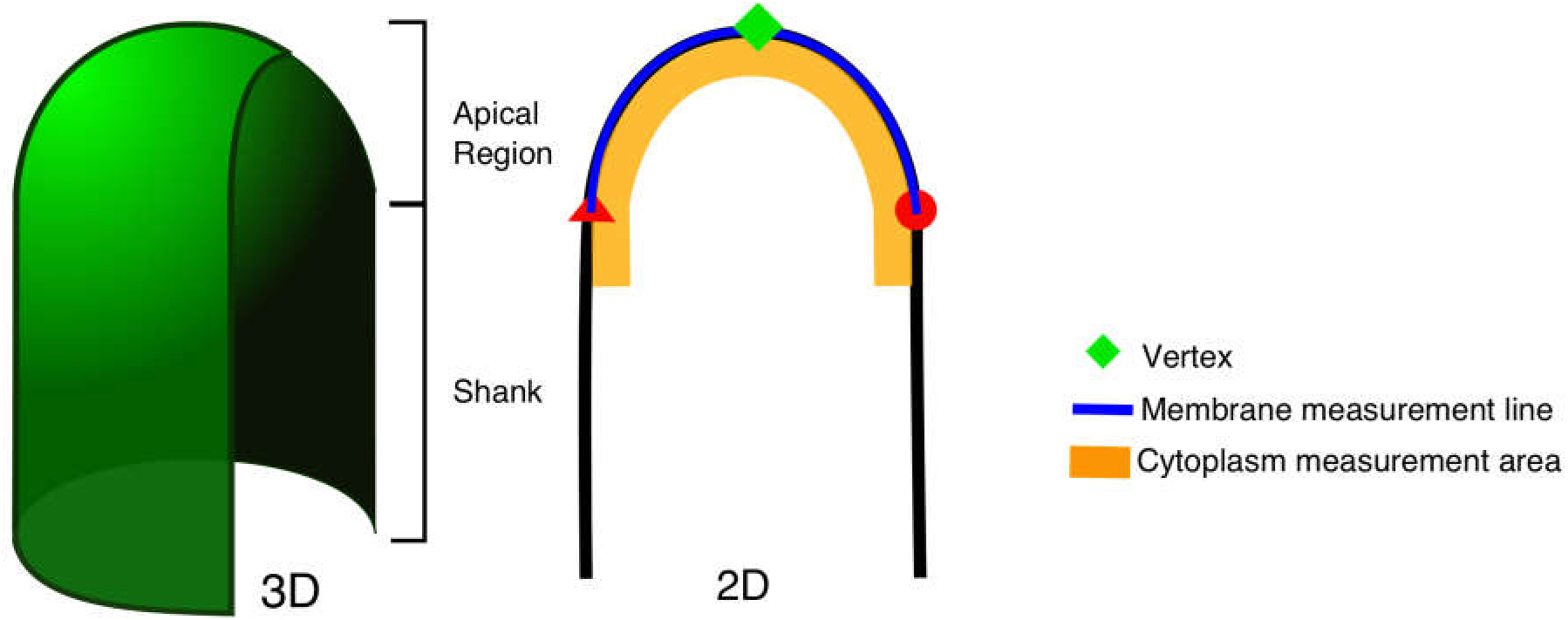
The schematic diagram of the apical region and shank of the pollen tube in 3D with an accompanying 2D diagram. The 2D diagram shows the measurement regions: membrane (blue) and tip cytoplasm(orange)

#### Segmentation of the tip-growing cell from fluorescent images

The input images for our computational model have to be fluorescence images that show the localization and intensity of an FP-tagged protein inside the cell. Image segmentation treats fluorescence images as an intensity map. After smoothing the image with a Gaussian filter, a Laplacian is used to detect regions of varying pixel intensities (regions with edges). This is done using a LoG-filter that takes the form:

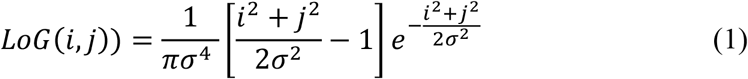

where *i* and *j* are the row and column positions of a pixel respectively, and 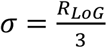 is the standard deviation of the Gaussian filter. An entropy filter (of radius *R_Entropy_*) is used to compute the entropy of the image. In this case, entropy is defined as the measure of contrast in a given region: regions with lower variation in pixel intensities have lower entropy as compared to regions with a larger pixel intensity variation. Otsu’s thresholding algorithm is used on the result of the entropy calculation to obtain a segmentation of the image. The initial values for *R_LoG_* and *R_Entropy_* are user-defined. At the start of the program, the user is allowed to change these values to ensure accurate segmentation of the cell region.

While this is a very effective segmentation procedure, it creates some false positive regions. Simple morphological operations (erosion (*radius* = 0.7*r_ent_*) followed by hole-filling) are used to obtain simply connected regions without holes. Regions with sizes of less than 10 pixels are discarded. Of the remaining regions, the region with the highest average intensity is designated as the cell region. The result of this process is a binary image that shows a single cell region as the foreground (pixel value = 1). This binary image is smoothed to promote contour smoothness. This technique has been used in our previous work segmentation. Shape smoothing is the final phase of the segmentation process, which is done to promote a smooth contour of the cell shape. A smooth contour is helpful in determining the curvature along the contour of the cell. The smoothing process is an iterative process performed on the binary segmentation, and it terminates when there is no change in the shape of the cell between consecutive iterations or the maximum number of iterations (500) is reached. Let the result of segmentation be the image *I* of size *r* × *c* (r-rows and c-columns). At each iteration, a temporary image 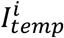 (temporary *I*mage at iteration *i*) is created as:

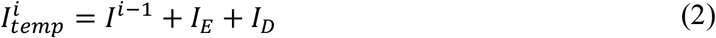

where *I*^*i*−1^ is the result of the previous iteration, *I_D_* is formed by adding a layer of pixels to the border of the object in *I*^*i*−1^ and *I_E_* is formed by removing one layer of pixels at the border of the object in *I*^*i*−1^. 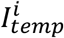 has values between [0 3]. This temporary image is filtered using:

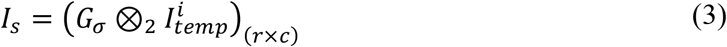

where ⊗_2_ is the 2D-convolution operator, *G_σ_* is a 2D-Gaussian filter with a standard deviation of *σ*. (·)_(*r* × *c*)_ enforces the condition that 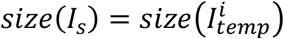. This is achieved by zero-padding the edges of 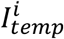 with 3*σ* rows/columns to each edge. In this version of the program, the standard deviation of the filter is set to 1.5 *pixels*. A new segmentation is created from the smoothed image by applying a threshold the following thresholding scheme: a pixel location (x, y) is considered stable (*I*(*x, y*) ≥ 2) if it belongs to two of the three sub-images in equation (1) namely *I*^*t*−1^, *I_E_* and *I_D_*. Applying this threshold creates a new binary image as:

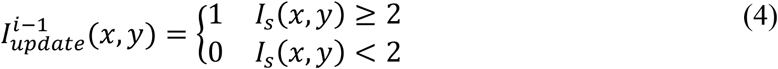

The new binary image 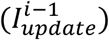 is compared to the result of the previous iteration (*I*^*i*−1^). If there is a change in the shape, then equations (2) - (4) are repeated. If there is no change in the shape of the object, a final image is produced by dilating 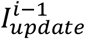 using a disk element:

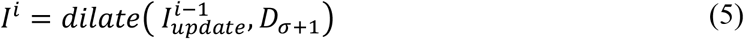

where *dilate*(·) is the dilation function, *D* is a disk element of radius *σ* + 1. *σ* has the same definition as in (**Error! Reference source not found.**).

#### Estimation of the apex of the tip-growing cell

After segmented the tip-growing cell from the given images, we aim to identify the apex (tip vertex) of the apical growing tip. Estimation of the apex of the growing tip is modeled as an ellipse that best describes this two-dimensional region. The conic section whose equation is given by:

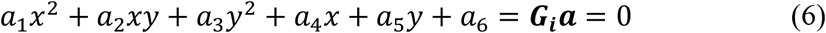

represents an ellipse^18^ if 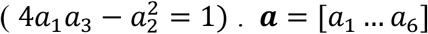 are estimated coefficients, ***G*** = [*x*^2^, *xy*, *y*^2^, *x*, *y*, 1] is an (*N* × 6) data matrix corresponding to the *i^th^* 2D-line segment and {(*x_i_*, *y_i_*), *i* ∈ 1, …, *N*} represents the x- and y-coordinates of the points on a line segment. The center of this ellipse (*c_x_*, *c_y_*) is obtained using:

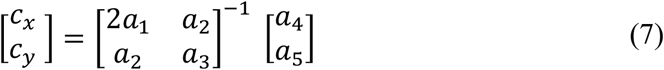

Given that the equation of an ellipse can be made to fit almost any continuous line segment of the cell shape, the task becomes one of finding the optimal/best location for the tip. The intuition for our solution is that overlapping line segments around the tip will have different coefficients (***a***_*i*_), but all of them will have approximately the same ellipse center (*c_x_, c_y_*). So, the tip candidates are the contour points whose ellipse center location has the most votes. In the ellipse-fitting process, the contour is split into overlapping segments of length:

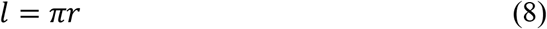

where *r* is the radius of the cell region. This radius (*r*) is based on the cell shape obtained from segmentation (Figure 2A(I)). A distance transform on this shape finds the minimum distance of each cell pixel to the cell contour. *r* is the maximum value of these distances.

**Figure 2:**
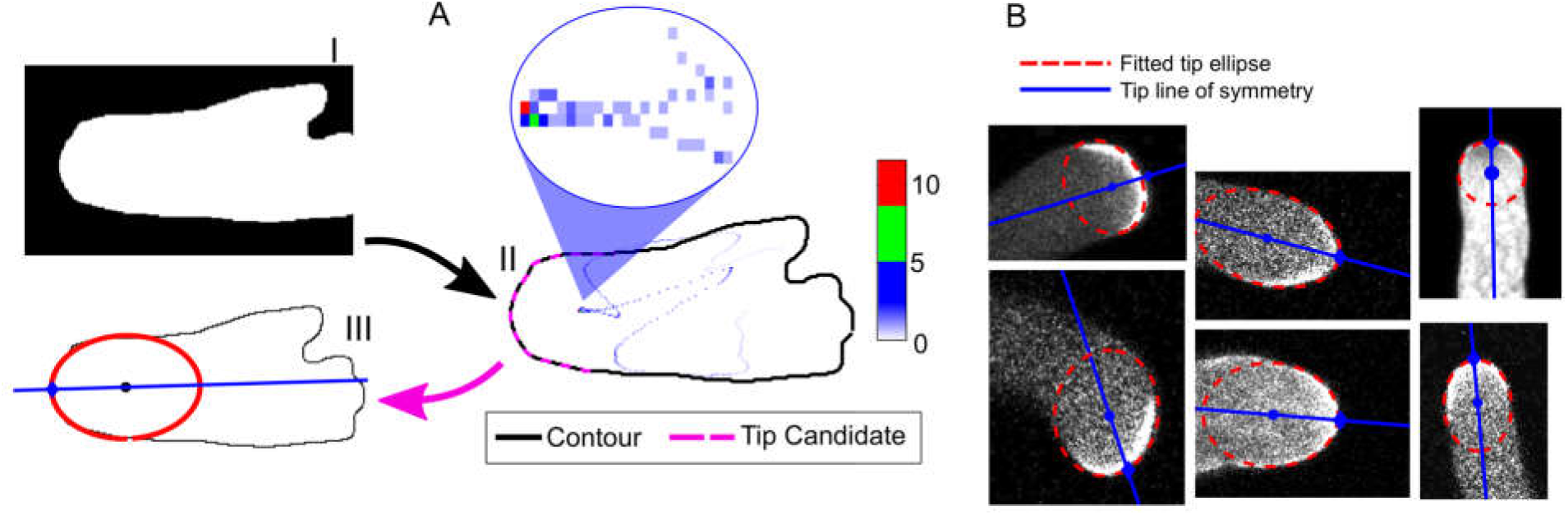
Tip location procedure on a sample-segmented image. (A) The tip localization method starting with the segmented image (I). Image (II) shows the cell contour, the tip candidates, and the votes received by each point as an ellipse center. A close-up of the tip region shows the center-location with the highest votes (in red). Image (III) shows an ellipse that is fitted to the tip candidates, and the vertex location. (B) Examples of tip detection on sample images with various tip widths. The solid line indicates the tip line of symmetry that is used to determine the tip vertex.

We use a voting scheme to search for the tip region. For each line segment (*l*) from the cell contour, the solution to equations (6)-(7) provides the location of the ellipse center^18,19^. The vote at this center location is incremented by 1. This voting process continues for all the line segments. The set of line segments, whose ellipse-center location gets the highest votes, become the candidates for the contour of the cell tip (see Figure 2 A(II) where the dotted points inside the cell indicate the number of votes received by that location as an ellipse center). The result of this process is the list of candidate tip locations.

Note that in the images shown in Figure 2B, the major axis (longest radius) of the tip ellipses is not guaranteed to fall at the correct vertex of the tip region (e.g. Figure 2B, top left and bottom center). The location of the vertex is important because it denotes the geometric center of the tip region, from which point the distance *arclength* (a user-controlled parameter that defines half the length of membrane measurement line) is determined for membrane measurements. To find the best (optimal) vertex location, we employ the following iterative scheme. After the tip detection phase, the current tip candidates (Figure 2A-II) enclose only a small portion of the cell shape. As such, we iteratively add one pixel (one point) of the cell contour to each end of the tip candidates, thereby including more of the shank in the tip region. An ellipse is fitted to this new region and the intersection between the major axis of this ellipse and the tip candidates is noted. This process of increasing the tip region continues until the last five intersection-locations between the tip candidates and fitted ellipses are the same. Alternatively, the process can go on for fifty iterations and the average of the tip intersection, weighted by the corresponding tip area, becomes the tip vertex location.

When analyzing a video sequence, we impose the condition that the pixel locations of the tip in the current frame should be close to those in the previous frame. As such, instead of repeating the entire procedure outlined above, we focus on the points in the new curve that are within a specified distance of the tip points in the previous frame. This distance is set to 25% of the detected cell radius at the tip. The newly acquired tip candidate points undergo the above curvature method for vertex detection and the *arclength*-parameter is used to find the tip points for the current frame.

#### Finding the optimal membrane measurement location

The result of segmentation is the contour of the cell. This contour represents where the cell wall/membrane of the cell. However, the location of this contour may not necessarily match with that drawn by someone with knowledge of the pollen tube shape. Resolving this discrepancy is important in ensuring that the membrane measurements made by TipQAD closely resemble those obtained by manually drawing on the tip region. This resolution requires a search, from the original tip contour in the direction of the cytoplasm, for the optimal measurement location. We incorporate a *winsize* parameter, which determines how far into the cytoplasm the algorithm should search for the best measurement line. If *winsize* = 0, then the original tip contour is used as the measurement line. Large values of *winsize* increase the search area for the membrane line. The value of *winsize* should be based on the radius of the cell: if the images have small pollen tubes, then *winsize* should be less than 5.

#### User-defined measurements of cytoplasm intensity through adjusting two parameters (*Tiplength, Depth*)

The measurement region within the cytoplasm is determined by two parameters: *Tiplength* and *depth*. *Tiplength* parameter is the length of the region along the cell contour on either side of the vertex. This means that the width of the measurement region from end-to-end is 2 × *Tiplength*. The *depth* parameter determines how far into the cytoplasm the measurement region goes. The values for these parameters can be altered from their default values to best match the desired measurement region. For a given cytoplasm region of an image, the average pixel value within the region is recorded. If *depth* = 0, then the measurement region is the cell contour at the tip region with a width of 2 × *Tiplength*.

Figure 3 shows the results of comparing measurements from automatic cytoplasm regions to manual measurements. The manual measurements were obtained using ImageJ software and a sample of the measurement region is shown in the first image of Figure 3(a). The rest of the images in Figure 3(a) show cytoplasm regions for *Tiplength* = 20*μm* and various values of *depth*. We explore different combinations of values for *tiplength* and *depth*:

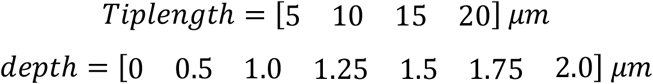

**Figure 3:**
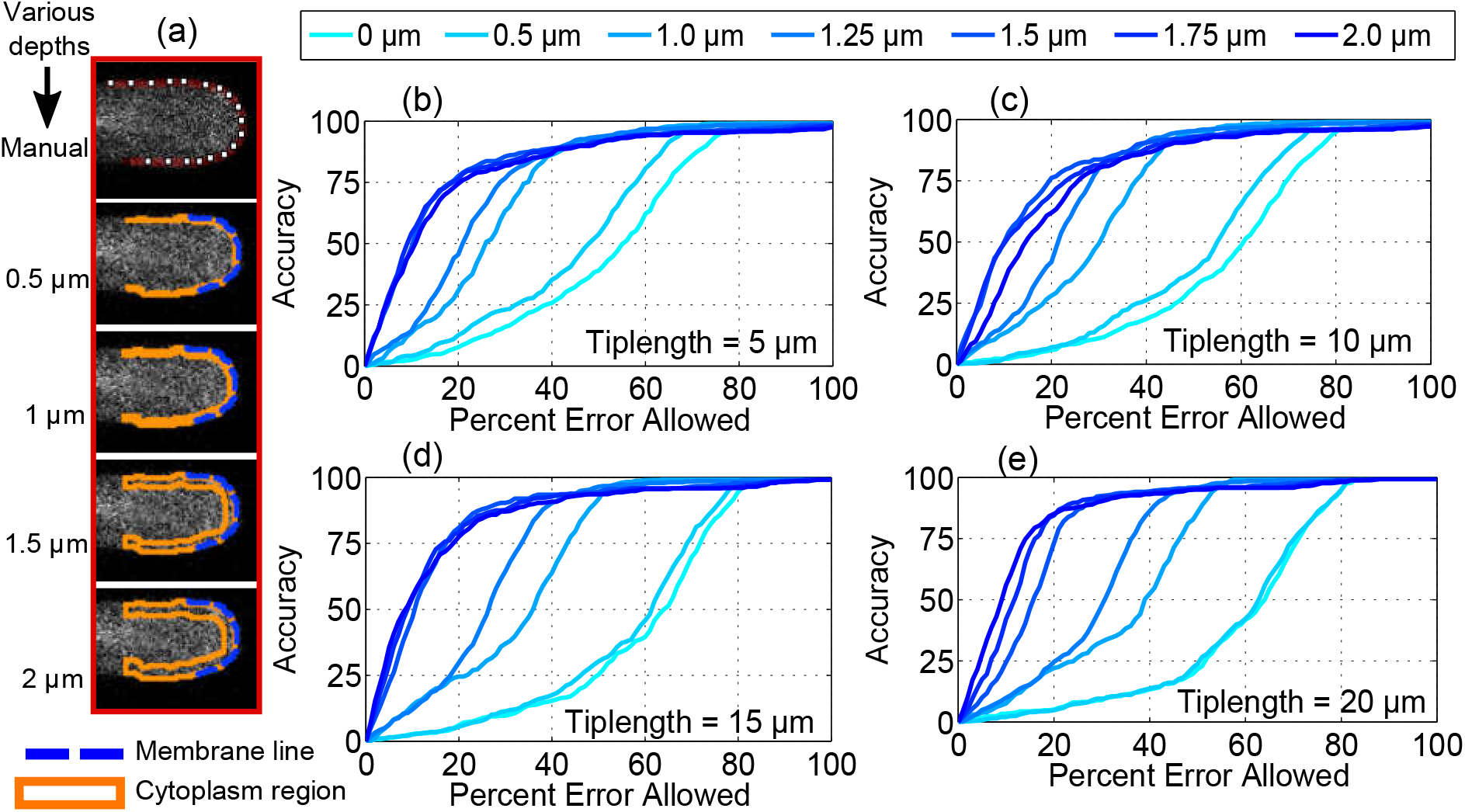
The effect of *Tiplength* and *Depth* parameters on tip cytoplasm measurements over four videos (826 images total). (a) Sample image showing measurement regions. The first image shows the manually drawn measurement region. The remaining 4 images show automatic cytoplasm measurement regions (orange) with increasing measurement *depths* when *Tiplength* = 20*μm*. (b)-(e) Median of cytoplasm measurement accuracy at specific values of *Tiplength* for various values of *depth* over 4 experimental videos. Plot (e) has the most accurate values when *Tiplength* = 20*μm* and *depth* = 1.75*μm* or 2*μm*. The measurement regions for these two depths are the last two images of (a) respectively.

The plots of Figure 3(b)-(e) show the results of using a given *tiplength* and all the *depth* values for the four videos used. The y-axis is the median accuracy and the x-axis is the tolerated percent error between the automatic and manual measurements (the threshold used to classify an automatic measurement as correct or incorrect). According to these plots, the most accurate *depth*-values are 1.75 *μm* and 2.0 *μm* regardless of the *tiplength*. However, the highest accuracies are achieved when *Tiplength* = 20*μm* and *depth* = [1.75 or 2]*μm*. Incidentally, these measurement parameters define a region that is similar in location and size to the manually drawn measurement region.

#### User-interface of our tool TipQAD

Figure 4 shows a snapshot of the TipQAD user interface. The interface is grouped into 3 major regions: segmentation parameters, measurement parameters, and analysis. At the start of the program, the user chooses the type of source files (video, folder, or .lif file). The default <u>Segmentation Parameters</u> can be altered to ensure that the segmentation captures the entire pollen tube. The default <u>Measurement Parameters</u> can also be changed and the measurement regions visualized to ensure proper measurements. Lastly, the <u>Analysis</u> parameters are set and the program analyzes the specified frames. At the end of the analysis, the user can visualize measurement regions and the associated tip membrane distribution by selecting a frame index from the “Display Regions on Image” menu. The accumulated data can be saved by clicking on “Save Data”. Table 1 lists the input parameters for TipQAD as well as a short description of each parameter and its default value. At the start of the program, the user can alter the values of these parameters.

**Table 1:** Table describing user inputs to TipQAD, their units and default values.

| Input | Description [*units*] | Default |
| --- | --- | --- |
| R<sub>LoG</sub> | Radius of the LoG filter [*pixels*] | 17 |
| R<sub>Entropy</sub> | Radius of the entropy filter [*pixels*] | 9 |
| Smooth contour | (on = yes, off = no) Smooth segmented contour | on |
| Resize images | (on = yes, off = no) Resize the image to $[256 \times 256]$ | on |
| Winsize | Width of fluorescence search space [$\mu m$] | 5 |
| Pixel scale | Size of a pixel [$\mu m$] | 0.0758 |
| Arclength | Half the length of membrane measurement line [$\mu m$] | 10 |
| Tiplength | Half the length of cytoplasm measurement area [$\mu m$] | 10 |
| Depth | Distance from contour into the cytoplasm [$\mu m$] | 2 |
| Mactime | Microscope acquisition time for a single image [$sec$] | 1 |
| Delay | Microscope delay time between two images [$sec$] | 1 |
| First frame index | Index of the first frame to be analyzed | 1 |
| Spacing | Spacing between analyzed images | 1 |
| Last frame index | Index of the last frame to be analyzed | Video length |
| Display results | (on = yes, off = no) Display dynamics during execution | on |

**Figure 4:**
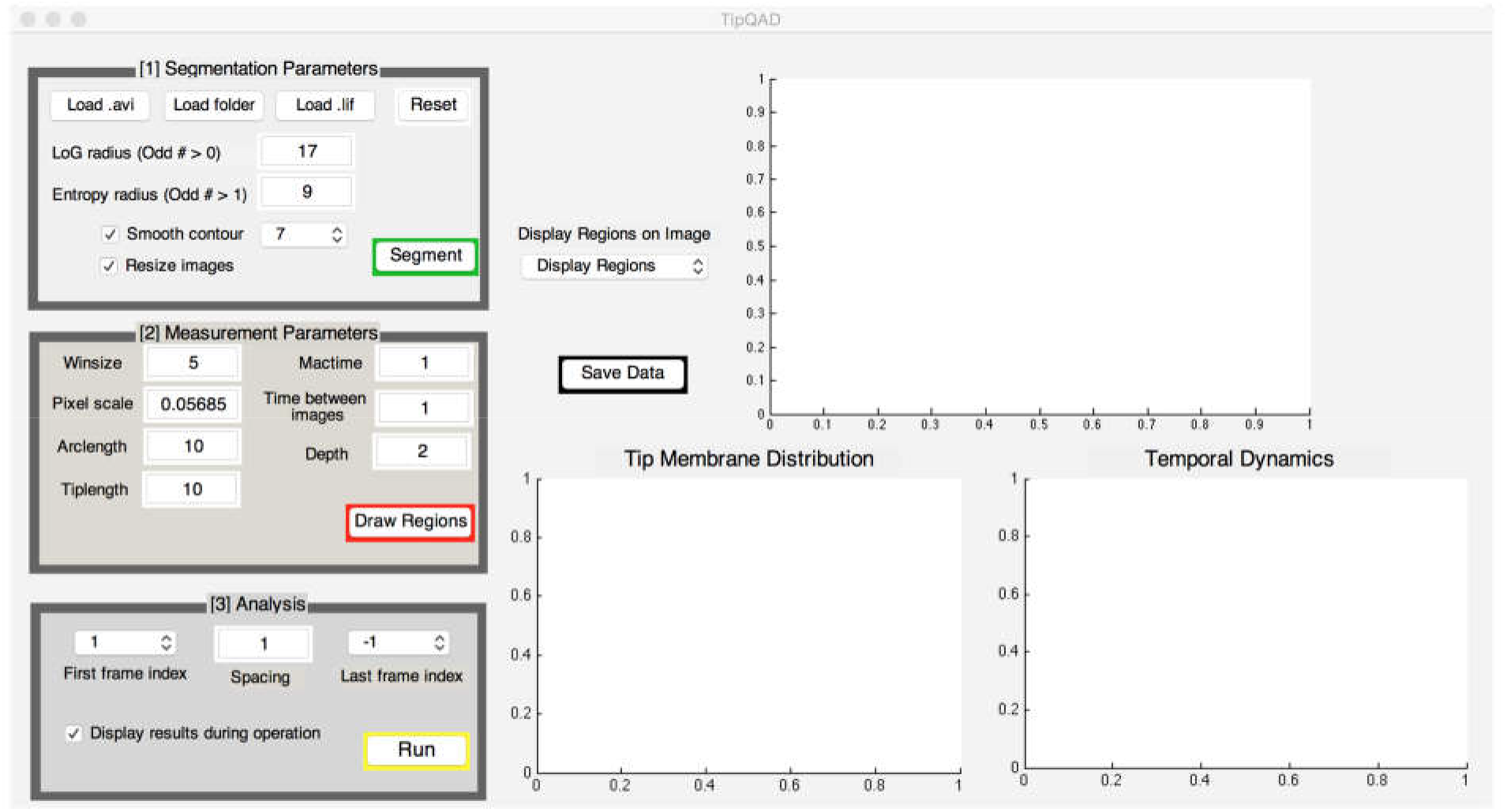
Display of cell analysis during program execution. TipQAD interface where the user selects what type of sequence to analyze, set/check segmentation parameters, set/check the measurement regions on the image, and set the range of images to be analyzed. As the analysis proceeds, the top image displays the previous image and measurement regions, the bottom left graph shows the fluorescence distribution along the tip membrane and the right graph shows the temporal average of both membrane (blue:solid) and tip cytoplasm (orange:dashed) regions. The “Display Regions on image” options is used after the analysis to visualize the measurement regions on any of the analyzed images. “Save Data” saves the analyzed data. By adjusting the First/Spacing/Last indices, the user can analyze (and save) selected portions of the dataset.

### Making measurements

In this section, we compare manual measurements for cell length, the intensity of PM protein fluorescence and actin filaments to those made by TipQAD. Manual measurements are obtained by trained users using ImageJ^20^ software. The accuracy of TipQAD is determined by the following equation:

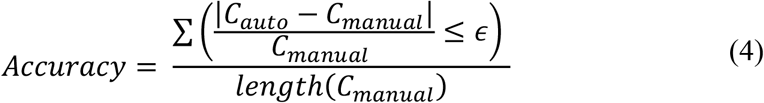

where |·| is the absolute value operator, *ϵ* ∈ [0 1] is the acceptance threshold (0 = 0%, 1 = 100%), *C*_(·)_ denotes the measurements obtained using TipQAD (C_auto_) and manual measurements (C_manual_). The numerator determines the number of measurement points that are within *ϵ* of the manual value, i.e. if an automatically measured value falls within *ϵ*-% of the manual value, then it is considered a correct measurement. For example, 90 is within 30% of 70 because 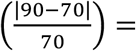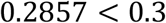.

### Cell Length vs. Time

The length of the tip-growing cell over time is determined by an analysis of the collected vertex locations. These locations form a trajectory of the tip in a video. To minimize fluctuations in this trajectory, we smooth the trajectory using a moving-average process. The smoothing process continues until the rate of change between two iterations, measured by the gradient of the mean-squared-error, falls below a specified threshold (1 × 10^−4^). The smoothed curve at iteration (t+1) is the result of a convolution operation between the curve at iteration (t) with a smoothing function. The Euclidean distance between the smoothed vertices is computed and summed to give the length of the cell over time. The cell length in the first frame is considered to be 0. Figure 5 shows length measurements and accuracy statistics for 5 videos of *in vitro*-growing *Arabidopsis* pollen tubes expressing CRIB4-GFP. The plot in Figure 5B shows that the accuracy of automated measurements increases as more error is accepted by the user. If the user accepts a 10% error, then on average, TipQAD is 75.5% close to the manual values. If the user accepts a 20% error, then on average TipQAD is 81.5% close to the manual values. The shaded region around the median line indicates the 1^st^ and 3^rd^ quartiles at each error threshold.

**Figure 5:**
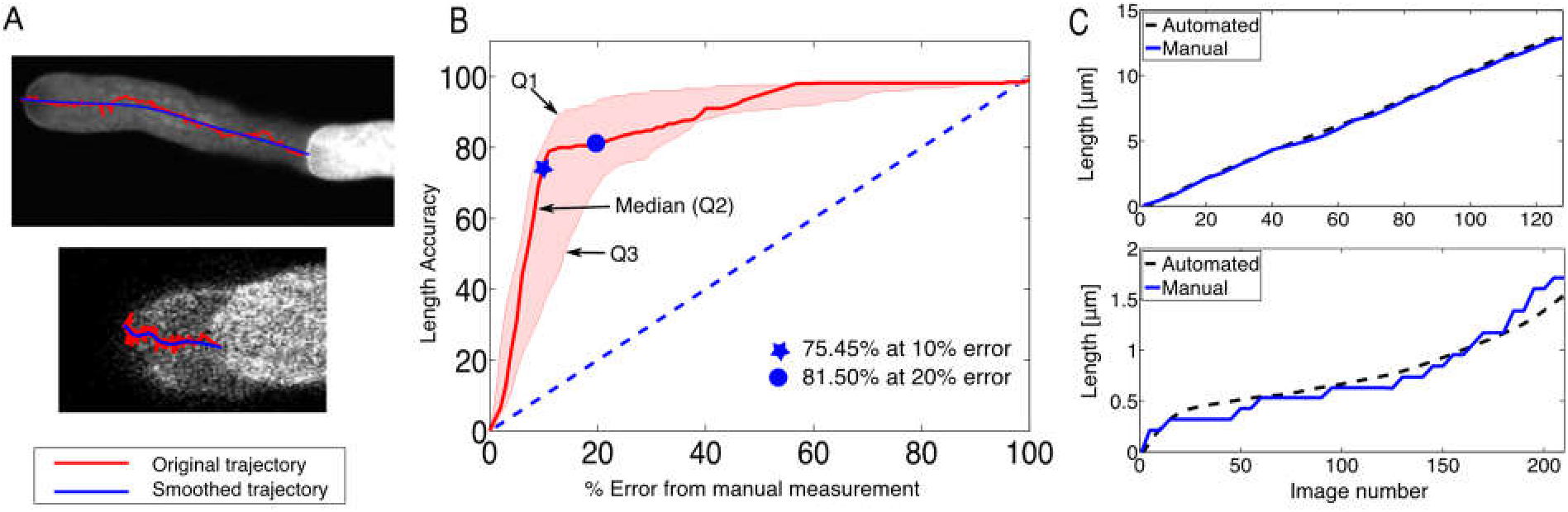
Comparing manual vs. automated measurements for cell length over time for 5 experimental videos. (A) Overlay of the trajectory of vertices on the start/end images from two videos. (B) Median value of accuracy, bounded by first and third quartile values, of automated length vs. error from manual length. (C) Examples of automated length measurements (dashed) vs. manual length measurements (solid) for two videos.

### Fluorescence Measurements on the PM of the Tip

In these experiments, we used GFP-tagged PRK1 (AT5G35390), which is a receptor-like kinase in *Arabidopsis* and localizes to the PM at the tip of pollen tubes, as a membrane marker protein. The model input parameters are at their default values except for the *arclength* whose value is set to 6*μm*. Membrane fluorescence measurements are made along the tip region. The desired measurement distance is specified by the user-defined variable *arclength*, which specifies a maximum distance on either side of the vertex along the PM. Contour (PM) points that are within the *arclength*-distance of the vertex are used for measurements. Since the contour is based on automated segmentation, there is no guarantee that it falls precisely at the optimal location (as drawn by a user). To alleviate this problem, we specify a distance of *winsize*-pixels from cell contour into the cytoplasm. The optimal measurement location for membrane dynamics is found in this region using seam carving^21,22^. Seam carving is a process of finding the minimum (or maximum) path of resistance, or total energy (seam) when traversing a region from one end to the other. For our purposes, seam carving finds the path of maximum fluorescence from one end of the tip measurement line to the other. Figure 6 shows a comparison of measurements made using ImageJ and the proposed method.

**Figure 6:**
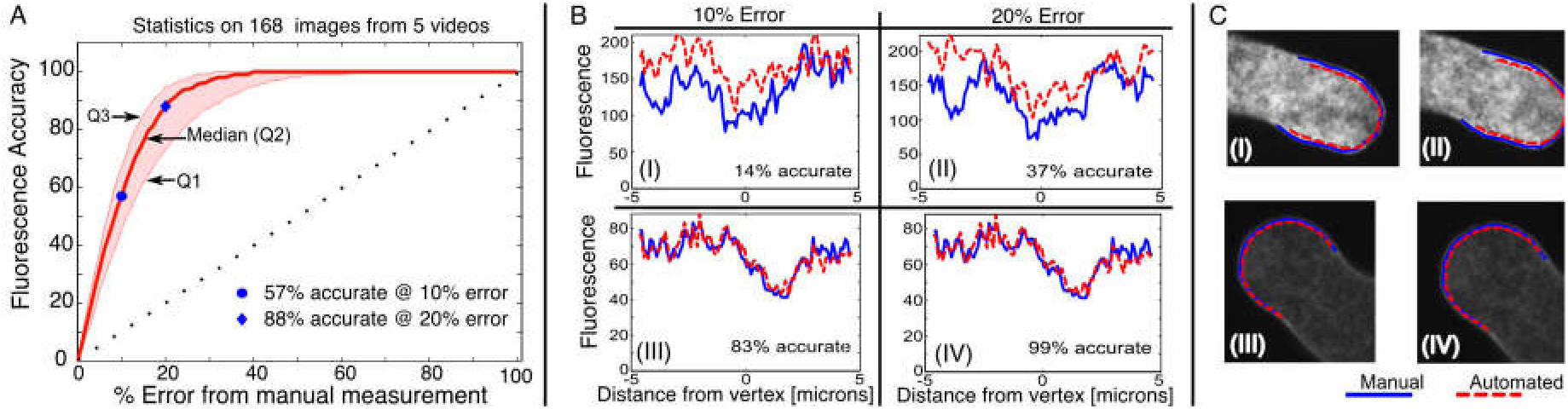
Comparing fluorescence measurements from user-drawn vs. model-detected regions. (A) Median fluorescence accuracy (bounded by first and third quartile statistics) for comparing automated measurements to manual measurements for 5 videos. (B) Sample tip fluorescence plots from manual-drawn (blue:solid) vs. model-detected (red:dashed) regions. At a threshold of 10% error, B(I) shows the most error and B(III) the least error. At a threshold of 20% error, B(II) shows the most error and B(IV) the least error. (C) Images showing manual and automated tip measurement lines. For each image, fluorescence values along measurement lines are shown in the corresponding image in (B).

### Fluorescence Measurements in the Tip Cytoplasm

Another important measurement in the study of tip growth is the distribution of cytoplasmic molecules or structures such as actin filaments, ions, and vesicles in the cell tip^23^. Figure 7A shows the selected regions for both manual and automated measurements using Lifeact-EGFP as a marker for actin filaments. It has been shown that both Arabidopsis and tobacco pollen tubes have a population of fine actin filaments in the cytoplasm close to the PM of the extreme tip^24^. The width and depth of the cytoplasm measurement region are user-defined parameters where the *width* = 2 × *tiplength.* In this region, we record the average fluorescence value and compare it with the results obtained by the automated process. Figure 8B shows the accuracy of automated measurements compared to the manual measurements. The x-axis indicates a measure of how far an automated measurement point is from the corresponding manual measurement (equation (4)). The y-axis shows the median accuracy value bounded by the first (lower) and third (upper) quartile statistics. This figure indicates that the automated measurements are close to the manual measurements.

**Figure 7:**
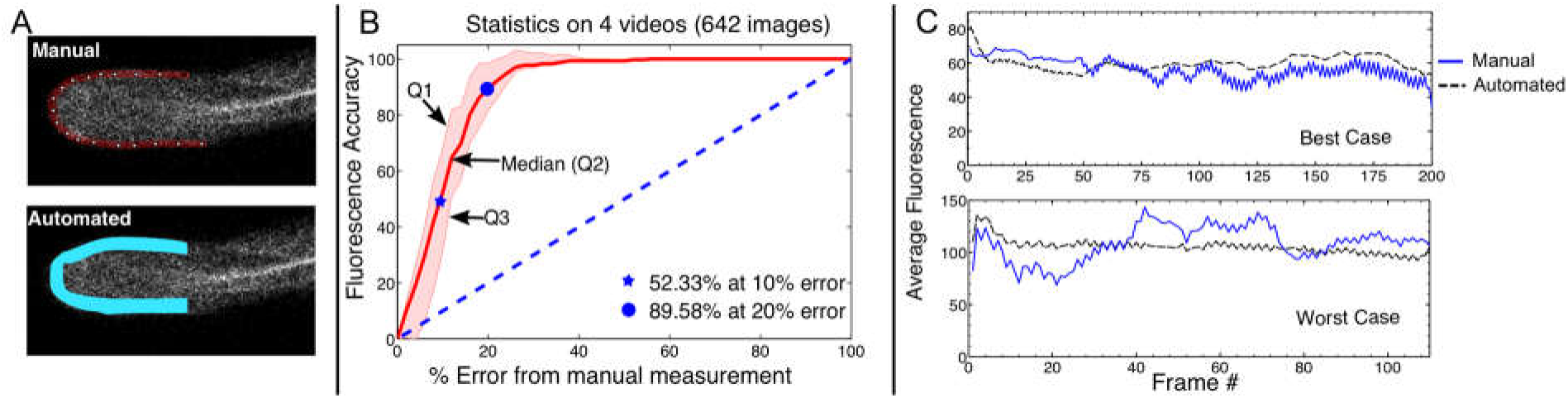
Comparing manual vs. automated tip cytoplasm measurements. (A) Sample images showing manual and automated measurement regions. (B) Plot of the median fluorescence accuracy bounded by first and third quartile statistics of the automated method. (C) Plots of manual and automated measurements of average fluorescence for 2 videos.

**Figure 8:**
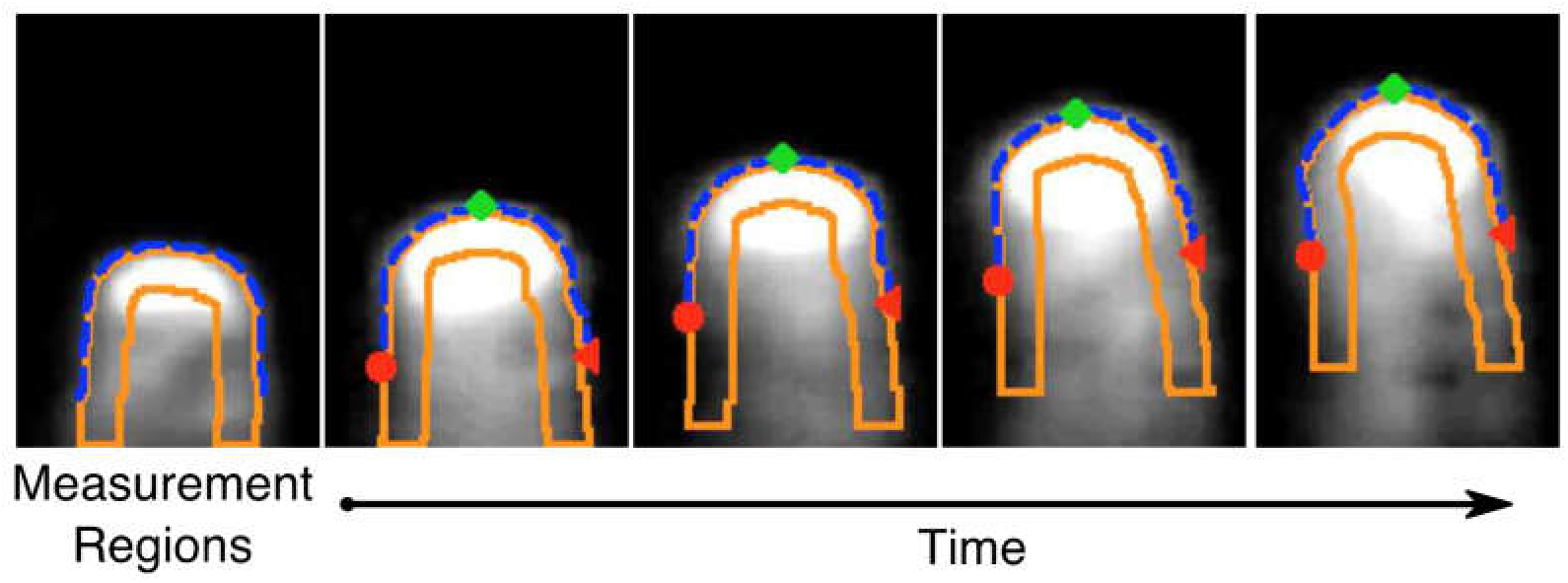
Sample frames from the analysis of a video of a growing root hair cell. The image on the left shows the initial measurement regions on the first frame of the video sequence. The subsequent images show the root hair at various growth stages with the associated measurement regions.

### Measurements on Root Hairs

As mentioned above, TipQAD can be used to analyze other tip-growing cells. Therefore, we took a previously published video of a growing root hair cell expressing YFP-RABA4b^25^ for analyzing using our tool, TipQAD. Figure 8 shows some sample frames from the above analysis. The detection and measurement processes are the same as described in the preceding sections. The first image shows the measurement regions overlaid on the first frame of the sequence. Subsequent images show selected frames and their associated measurement regions. In this example, the segmentation boundary is not as accurate as desired (e.g. on the fourth image). These inaccuracies are largely due to intensity variations in the image over time. However, TipQAD is able to obtain an estimate of the measurement regions relatively close to the expected locations.

### Application of TipQAD to FRAP (Fluorescence Recovery After Photobleaching) Analysis in Tip-growing Cells

TipQAD also has the potential to aid in analyzing FRAP experiments at the apical plasma membrane and in the cytoplasm. Figure 9 shows temporal measurements of fluorescence intensity in the tip region during a FRAP experiment, together with sample images at various time points. The video contains 24 frames of a FRAP experiment on a pollen tube expressing PRK1-GFP, where the PRK1-GFP on the tip PM was photobleached. The images are of size 512×512 and each pixel is of size 0.0758 *μm/pixel*. The parameters *arclength* and *tiplength* are both set to 6 *μm*. All other parameters are at their default values. The intensity in the membrane region is always higher than in the cytoplasm region. This is expected because the membrane measurement line follows the path of the highest intensity in the membrane region. Both measurement regions show a recovery of the fluorescence signal over time and a reduction towards the end of the sequence.

**Figure 9:**
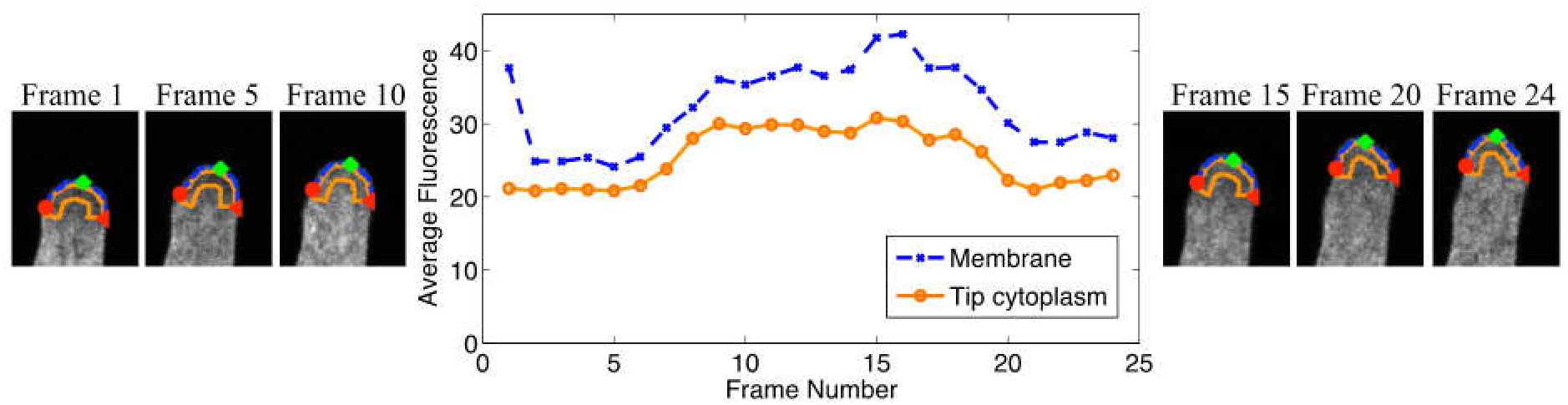
Plot of average membrane and cytoplasm fluorescence signals for a video sequence of a FRAP experiment on a pollen tube expressing PRK1-GFP, and some sample frames showing the measurement regions. The plots show a recovery of fluorescence in the tip region after photobleaching. The parameters *arclength* and *tiplength* are both set to 6 *μm*. All other parameters are at their default values.

## Discussion

TipQAD is a tool for quantifying spatiotemporal dynamics of molecules and structures in the apical region of tip-growing cells. The effectiveness of detecting the tip region relies on the cylindrical geometry of these cells. TipQAD allows automated analysis of video images in a given region of these growing cylindrical cells, which would otherwise be conducted in a manual manner using other software packages such as ImageJ. The novelty of TipQAD lies in the application of computer vision techniques to the quantitative analysis of dynamic molecular and structural behaviors in living cells. The cell segmentation process used in TipQAD relies heavily on the edge detection of fluorescence images throughout the growing region of the cell. If the fluorescent signal is only presented in the tip region, the detection process is likely to fail. However, this dependence can be overcome by implementing a segmentation method that fuses both fluorescence and bright field image modalities^22^.

TipQAD greatly reduces the time for analyzing and quantifying molecular and cellular dynamics in tip-growing cells. For a set of 1000 images, average processing times for TipQAD vs. manual analysis are approximately 15 *minutes* vs. 250 *minutes*. Temporal constraints on detecting the tip region ensure consistency in tip localization over time. Furthermore, automated analysis eliminates problems posed by human error and subjectivity, hence, it gives unbiased results. With this tool, hours of user-assisted analysis are reduced to a fraction of that time. This will enable biologists to quickly analyze experimental videos and identify trends in fluorescence signals from large amounts of video data.

We also demonstrated that TipQAD is able to analyze the FRAP experiment. Since the basis of FRAP and FRET data analysis is the measurement of PM or cytoplasmic fluorescence signal, our method can be directly applied in FRET experiments as well. In FRAP, the fluorescence intensity at the photobleached membrane regions can be readily measured with our method after adjusting the value of *arclength*, and the data can be used to calculate the recovery rate. In FRET, if the protein-of-interest is localized on the cell membrane or the apical region of the cytoplasm, our method can be used to measure the fluorescence intensity in the donor, the acceptor, and the FRET channel images. The data will then be processed in the standard procedure to calculate the bleedthrough ratio, the net FRET signal, and the FRET index. In both FRAP and FRET experiments, there is usually a large number of images or videos that need quantification. Our automated tool will be very useful in these applications for its high-throughput capability.

## Methods

### Plant Materials

The PRK1-GFP fragment was cloned from pLAT52∷RLK-GFP described in Lee et al., 2008, and was inserted into a binary vector, pCL, which was constructed by inserting the pollen tube specific promoter LAT52 into pCAMBIA1300 using SalI and XbaI restriction sites. The resulting vector pCL∷PRK1-GFP was introduced to wild-type Arabidopsis thaliana Col-0 plants using Agrobacterium-mediated floral-dip method.

The *Arabidopsis* plant expressing Lifeact-EGFP (pCAMBIA1300-Lat52-Lifeact-EGFP) seeds were provided by Prof. Shanjin Huang (Chinese Academy of Sciences, Beijing) ^26^. The level of expression was determined using ImageJ by drawing a line of 5-pixel width and 20 μM in length around the pollen tube tip and as close to the edge of the pollen tube as possible. The mean intensity of LifeACT-GFP along this line was then calculated.

### Plant Growth and Pollen Tube Preparation

Transgenic *Arabidopsis thaliana* plants were grown at 22°C in a growth room under a light regime of 16h of light and 8 h of dark. To generate pollen tubes for imaging, pollen grains were germinated at 28°C for 2-3 hours on a solid pollen germination medium containing 18% (w/v) sucrose, 0.01% (w/v) boric acid, 1 mM CaCl_2_, 1 mM Ca(NO_3_)_2_, 1 mM MgSO_4_, and 0.5% (w/v) agar.

### Confocal Imaging

Imaging of PRK1-GFP and Lifeact-EGFP in pollen tubes was performed on Leica SP2 and SP5 laser scanning confocal microscopes, respectively. PRK1-GFP and Lifeact-EGFP were excited with a 488 nm Argon laser line and detected at 498-560 nm. For the FRAP experiment on the pollen tube expressing PRK1-GFP, a 405 nm laser was used to photobleach the PRK1-GFP on the apical PM as described in Lee et al., 2008.

### Manual Image Quantification

Manual image quantification was performed using the ImageJ software (version 1.45s). To measure the fluorescence intensity on the plasma membrane, a segmented line was drawn along the cell periphery at the tip region. To measure the fluorescence intensity in the cytoplasm, a polygon was drawn to cover the region of interest.

## Acknowledgements

This work is funded by NSF Grant *DGE 0903667* to Dr. Bhanu as the lead PI and Tambo (as NSF IGERT Fellow), and National Institute of General Medical Sciences grant (*R01GM100130*) to Dr. Zhenbiao Yang.

## Author Contribution

A.L.T. and B.B. designed and implemented the computational method, performed data analysis and contributed to the paper. N.L. performed data collection and contributed to the paper. D.R., C.C., and I.L. performed data collection and contributed to the paper. F.W. contributed to the computational method. Z.Y. designed the experiments, supervised the data collection and contributed to the paper. All authors participated in various discussions.

## Additional Information

### Competing Financial Interest

The authors declare no competing financial interest.

